# Human gut metagenomes encode diverse GH156 sialidases

**DOI:** 10.1101/2022.06.28.497753

**Authors:** Evan Mann, Shahrokh Shekarriz, Michael G. Surette

## Abstract

The intestinal lining is protected by a mucous barrier composed predominantly of complex carbohydrates. Gut microbes employ an array of glycoside hydrolases (GHs) to liberate mucosal sugars as a nutrient source to facilitate host colonization. Intensive catabolism of mucosal glycans, however, may contribute to barrier erosion, pathogen encroachment and inflammation.

Sialic acid is an acidic sugar featured at terminal positions of host glycans. Characterized sialidases from the microbiome belong to the GH33 family, according to CAZy (**<u>C</u>**arbohydrate **<u>A</u>**ctive en**<u>Zy</u>**me) database classification. A 2018 functional metagenomics screen using thermal spring DNA uncovered the founding member of the GH156 sialidase family, which lacks homology to GH33 sialidases and could not be taxonomically assigned. Subsequent structural analysis revealed critical active site residues. We sought to determine if GH156 sialidases are present in the human gut microbiome where they might contribute to mucous erosion.

A subset of GH156 sequences from the CAZy database containing key sialidase residues was used to build a Hidden Markov Model. HMMsearch against public databases revealed ∼10X more putative GH156 sialidases than currently recognized by CAZy. Represented phyla include Bacteroidota, Verrucomicrobiota and Firmicutes_A from human microbiomes, all of which play notable roles in carbohydrate fermentation. Genomic analyses suggested that taxa containing GH156-encoding genes may utilize host-glycans. Analyses of metagenomic datasets revealed that GH156s are frequently encoded in metagenomes, with a greater variety and abundance of GH156 genes observed in traditional hunter-gatherer or agriculturalist societies than in industrialized societies, particularly relative to individuals with IBD. A GH156 gene frequently detected in traditional populations was cloned from stool sample DNA and the recombinant protein exhibited sialidase activity with a fluorogenic substrate.

**Importance:** Sialic acids occupy terminal positions of human glycans where they act as receptors for microbes, toxins and immune signaling molecules. Microbial enzymes that remove sialic acids, sialidases, are abundant in the human microbiome where they may contribute to shaping the microbiota community structure or contribute to pathology. Furthermore, sialidases have proven to hold therapeutic potential for cancer therapy. Here we examined the sequence space of a sialidase family of enzymes, GH156, previously unknown to the human gut environment. Our analyses suggest that human populations with disparate dietary practices harbour distinct varieties and abundances of GH156-encoding genes. Furthermore, we demonstrate the sialidase activity of a gut derived GH156. These results expand the diversity of sialidases that may contribute to host glycan degradation and these sequences may have biotechnological or clinical utility.

## Introduction

The human colon houses trillions of bacterial cells from hundreds-to-thousands of distinct species representing several diverse phyla (1). This consortium, the microbiota, is critical for host immune system development, nutrient acquisition and colonization resistance, but has also been implicated in the onset of non-infectious chronic conditions (2, 3). Inflammatory bowel disease (IBD), including Crohn’s disease (CD) and ulcerative colitis (UC), is one such condition which is increasing in incidence and prevalence globally, particularly in post-industrialized societies (4). IBD is characterized as an uncontrolled and vigorous immune response to the gut microbiota which can occur in predisposed individuals. However, the direct roles that the microbiota play in IBD etiology have yet to be fully resolved (5, 6), and it has become apparent that the host glycome is a critical factor in IBD onset and progression (7–10).

Collectively, N- and O-glycans on host proteins play myriad roles, notably in cell-to-cell signalling, host-microbe interactions and immune system regulation (11) and the enzymes that contribute to their destruction are recognized virulence factors (12–15). The colonic epithelium is protected from direct assault by a glycocalyx overlaid by a carbohydrate-rich mucous bilayer embedded with antimicrobial proteins (16). The mucous barrier is critical to gut health as evidenced by the observation that *Muc2*^-/-^ and Muc2-glycosylation impaired mice develop colitis spontaneously (17–20). While the inner mucous layer is sterile (21), the outer mucous layer is an extensively colonized microbial niche (16, 22, 23). Critically, the mucous layer is often thinner during inflammation, and bacteria encroach towards the epithelial surface where they are better positioned to induce an immune response (24, 25), akin to mice with genetically compromised mucous barriers.

Host glycans, composing the bulk of the mucous, offer a plentiful nutrient source for the microbes capable of depolymerizing them into constituent sugars (26–29), as well as those that scavenge the mucosal sugars released into the environment. Overall, host-glycan consumption is essential to the maintenance and persistence of a stable microbiota capable of providing colonization resistance to enteric pathogens, and its been reported that mucosal glycan structures, at least insofar as blood group antigen type and secretor status are concerned, can help select microbial community composition (30–35). However, the repertoire of glycans produced by the host (and accessible to the gut microbiota) is altered in response to inflammatory signals (36, 37), which act as complementary immunoregulatory signals (7, 8). These altered glycoforms effectively shift the mucosal nutrient availability, which may in turn alter the microbiota community structure. The rate of host-glycan turnover could be important for mucous layer integrity, with over-consumption of mucin glycans contributing to inflammation-associated barrier erosion. The microbiota of mice fed fibre-deficient diets increasingly relies on host glycans (38, 39), which compromises the mucous barrier and sensitizes the host to colitis (25, 40).

Bacterial catabolism of glycans occurs in a coordinated stepwise manner, with individual glycoside hydrolases (GHs) sequentially removing specific monosaccharides or oligosaccharides that can be completely metabolised within the cell (27, 41–43). GHs are classified within the CAZy database (http://www.cazy.org/) based on primary sequence similarity (44–47). There are presently >165GH families defined in the CAZy database, with each member of a given family possessing a conserved three-dimensional fold and set of catalytic residues. Family membership does not strictly define substrate-specificity (in terms of sugar and linkage types), but in many families reported activities are limited, depending on the size and sequence diversity of the family. Enzymes from several GH families have been demonstrated to contribute to host-glycan breakdown (27, 41, 48–51), and new N- and O-glycan-targeting enzymes are being actively reported (52–55). Together, these factors hinder bioinformatic annotation of a given GH’s macromolecular target and a microbe’s carbohydrate preferences. However, grouping closely related family members into subfamilies, clades, or clusters provides more robust substrate predictions for distinct ‘monofunctional groups’ (56–62).

Sialic acids occupy terminal positions of vertebrate N- and O-glycans predominantly via α2-3 and α2-6 linkages and they are often the first components removed by microbiota GHs before the glycan core can be accessed (48). *Sialic acid* refers to a group of 9-carbon sugars characterized by a C1 carboxylic acid, an exocyclic 3-carbon sidechain, and, often, (acetyl) modified amino and hydroxyl groups. The variety of distinguishing epitopes make sialic acids ideal for information transfer, notably in immune signaling processes (eg. by Siglecs), or use as receptors for enteric pathogens and toxins. The predominant sialic acid on human glycans is 5-(acetylamino)-3,5-dideoxy-D-glycero-α-D-galacto-non-2-ulopyranosonic acid (Neu5Ac) (63). Sialidases, the GHs that remove sialic acids, are important for colonization and are often considered virulence factors. Free sialic acid, released into the environment by commensals, is implicated in enteropathogen expansion due to Neu5Ac foraging (64–66). To date, the only gut bacterial sialidases implicated in N- and O-glycan breakdown belong to the CAZy family GH33 (48, 67, 68). GH33 enzymes display a six blade β-propeller fold, and the active site includes a trio of Arg residues stabilizing the Neu5Ac carboxylate group, a Glu/Tyr/Asp catalytic triad and a hydrophobic pocket to accommodate the amino-linked (acetyl) sidechain at position 5.

The sialidase family GH156 was established in 2018 (69). The founding member of the family, subsequently designated EnvSia156, was identified with a functional metagenomics approach, using DNA isolated from a thermal spring that could not be traced back to a known taxon. This enzyme displayed α2-3 and α2-6 Neu5Ac and Neu5Gc sialidase activity on a variety of substrates, including glycoproteins, glycolipids and sialolactose. Interestingly, the GH156 catalyzes an inverting reaction, while family GH33 strictly contains retaining enzymes. Successive work revealed that the EnvSia156 displays a (β/α)_8_ barrel fold, distinct from the six blade β- propeller fold of GH33 family-members, further illustrating the absence of shared phylogeny (70). Cocrystalization of the enzyme in complex with sialic acids strongly suggested candidates for catalytic residues. Here, we investigated the hypothesis that GH156s from human gut metagenomes contribute to the degradation of host sialosides.

## Results

### Construction of a GH156 sialidase profile Hidden Markov Model

A profile Hidden Markov Model (pHMM) for the GH156 family was created to enable search of publicly available protein databases. To facilitate construction of the model, we leveraged the available structural information for EnvSia156 as well as the GH156 sequences catalogued in the CAZy database (44–46). The proposed catalytic site, based on co-crystal structures with products and substrate analogs, includes a catalytic D14-H134 dyad as well as a conserved pair of Arg (R129 and R202) and an Asn (N346) positioned to stabilize the carboxylate group (72). The 51 GH156 sequences in the CAZy database (March 2020) were aligned and those sequences lacking the catalytic Asp-His dyad or carboxylate-coordinating Arg-Arg-Asn triad, or suitable (i.e. polar) substitutions, were pruned from the alignment. We reasoned that this would favour the identification of homologs that hydrolyze sugars with a C1 carboxylate group, such as sialic acids, over potential GH156 family members that may target neutral sugars. The 32 CAZy sequences that met these criteria were truncated to the boundaries of the EnvSia156 (ß/α)_8_-barrel domain. HmmerBuild was run using the pruned and truncated multiple sequence alignment to construct the pHMM (Fig. 1A) (71). A sequence logo of the pHMM depicts the high weight of the catalytic and carboxylate-binding positions in the model.

**Figure 1.**
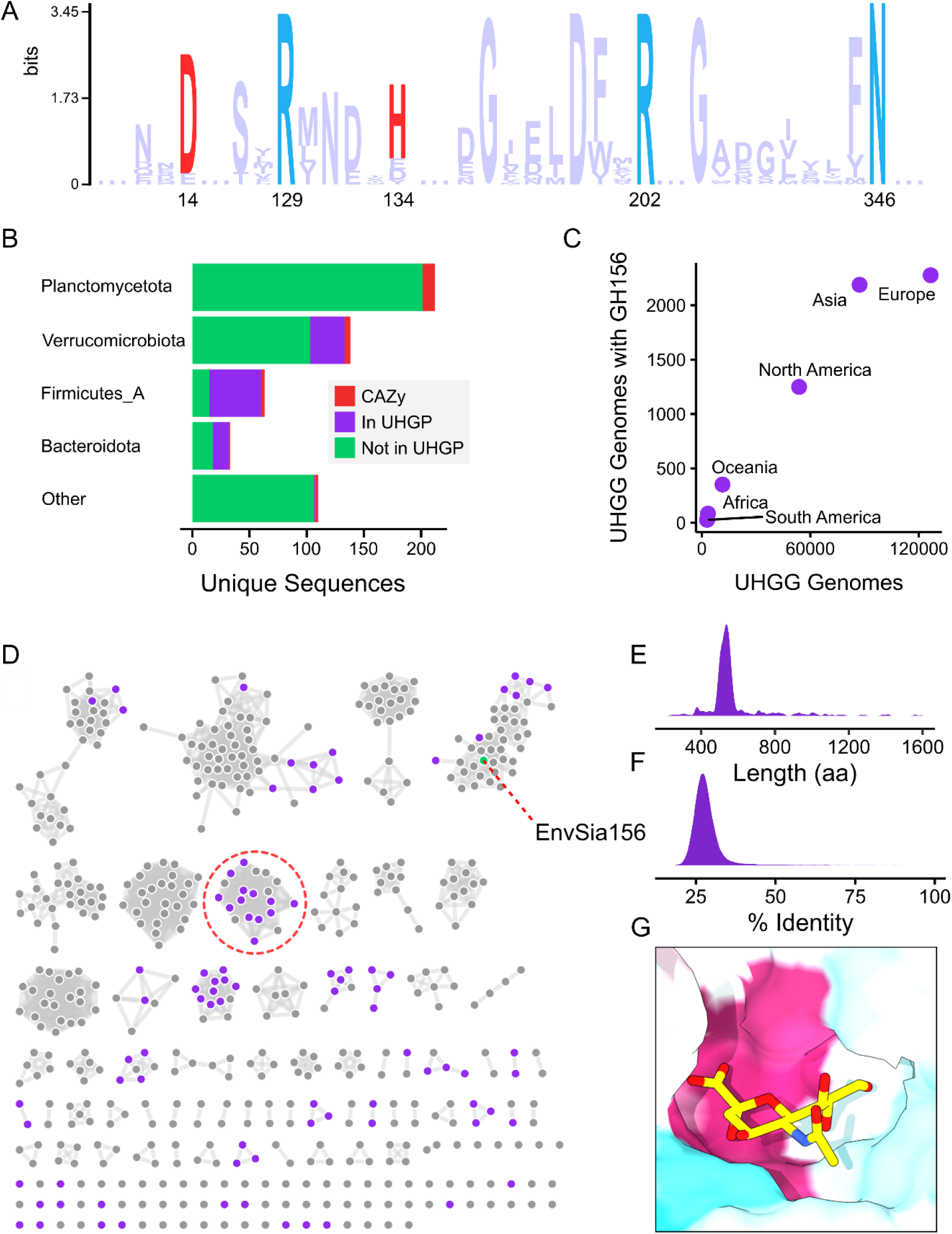
Diverse GH156s from distinct phyla were identified in global human gut metagenomes. A sequence logo of the HMM used for database searches denoting the relative importance of proposed catalytic residues (red) and carboxylate-binding residues (blue), numbered based on the EnvSia156 sequence (A). Taxonomic distribution of non-redundant GH156 proteins identified, highlighting the number of unique sequences from the human gut protein catalog (B). Regional distribution of GH156-encoding UHGG MAGs, demonstrating that GH156s are globally distributed (C). Sequence Similarity Network of non-redundant GH156 protein domain sequences (nodes) connected by edges indicating sequence identity >45%; nodes in purple indicate the sequence is derived from the UHGP-95 catalog (D). Nodes clustered in the red circle are derived from commonly identified Bacteroidota genomes (Table S3). Histogram of GH156 protein lengths (E). Histogram of catalytic domain pair-wise identity of GH156 family members (F). Sequence conservation mapped to the surface of EnvSia156 (pdb 6S00) in complex with Neu5Ac (yellow sticks) showing high conservation (magenta) around the anomeric carbon and carboxylate, and low conservation (cyan) around the exocyclic acetamido and glycerol groups.

### Identification of GH156s in protein databases

HmmerSearch was used to search three large public databases: UniprotKB’s over 225 million protein sequences (72), the Genomes from Earth’s Microbiomes protein catalog (GEM) consisting of over 5.7 million protein sequences derived from environmental metagenome sequencing projects (73), and the Unified Human Gastrointestinal Protein catalog (UHGP-95), consisting of 20.2 million non-redundant protein sequences from global human gut metagenome samples (74). Hits were manually inspected to ensure the presence of at least four of the five highlighted active site residues (accepting the substitution of a single one of these for a different polar amino acid). The hits were aggregated, and redundancy was eliminated using CD-HIT (75, 76) with default parameters and a 95% identity threshold. The resulting 556 non-redundant putative GH156 sialidase sequences were trimmed to the domain boundaries based on the pHMM alignment for subsequent bioinformatic analyses.

GH156s were identified across diverse taxa, with 18 phyla represented. The most frequent phylum was Planctomycetes with 212 GH156 sequences exclusively from environmental sources (Fig. 1B). GH156s from the UHGP-95 catalog represent enzymes that may be active against human sialosides in the gastrointestinal tract. Ninety sequences were derived from the UHGP-95 catalog: 45 from Firmicutes_A, 30 from Verrucomicrobiota and 14 from Bacteroidota species, as well as a single Proteobacterium (*Methylobacterium methylobacterium*). The only Verrucomicrobiota that could be assigned at the species level was *Victivallis vadensis*. Likewise, a single Firmicutes was assigned at the species level: *Mitsuokella jalaludinii*. This reflects that this family of enzymes in the human microbiome is predominantly found in poorly studied, low abundance or rare taxa. Globally, over half of the GH156-containing metagenome-assembled genomes (MAGs) from UHGG were assigned to *Parabacteroides merdae*, including 75% of those from North America.

Analysis of the geographic distribution of GH156-containing MAGs from the UHGG suggests that GH156s are globally distributed (Fig. 1C). While assembled GH156-containing MAGs are primarily from Asia, Europe, and North America, less sampled continents are the source of a roughly proportional number of GH156-containing MAGs, relative to the total number of MAGs that were assembled from a given region. Its likely that additional sampling of these under-studied regions will reveal further GH156 diversity.

To visually evaluate GH156 sequence diversity, we generated a Sequence Similarity Network (SSN) using the SSN tool developed by the Enzyme Function Initiative (77–80). We generated SSNs with edge thresholds from 40-60% identity (alignment length > 100 aa) in 5% increments. The maximum number of clusters with at least 10 sequences was at a 45% identity threshold, where there are 12 clusters with at least 10 sequences (Fig. 1D) and clustering was not strictly along taxonomic lines (Fig. S1A). At >60% identity, an approximate benchmark for intracluster substrate uniformity for glycoside hydrolases (56, 57, 81), there are only four clusters with at least 10 sequences, and a preponderance of singletons, doublets and triplets. Relatively few UHGP-95 sequences are in the same cluster as EnvSia156, providing limited support for a conserved function across the entire family. Collectively, the GH156s identified here share low sequence similarity, with a mean pairwise sequence identity (minimum 100 aa alignment) of 28.2% (Fig. 1F). This degree of sequence heterogeneity likely reflects a combination of incomplete sampling of this GH family but may also indicate that GH156s recognize diverse substrates. This is supported by sequence variability in the substrate binding pocket around sialic acid N-Ac and C7-C9 functional groups (Fig. 1G). However, the majority of Bacteriodetes sequences formed a single cluster with high sequence similarity (Fig. 1D, red circle) and these likely share a function across the different species.

### GH156 enzymes are modular

EnvSia165 possesses a C-terminal immunoglobulin-like domain, similar to some carbohydrate-binding modules, suggesting it likely plays a binding role. Indeed, most GH156s identified here were between 500-600 aa (Fig. 1E), consistent with a multidomain architecture with a carbohydrate-binding module. A subset of sequences were larger than 800 aa and we therefore predicted that they might contain an additional enzymatic domain(s). The separate domains in modular GHs often act upon the same glycan. GH156s active against human sialosides would be expected to be associated with other modules with activities targeting linkages seen in human glycans. We therefore annotated the identified GH156s using dbCAN2 (82) which revealed several distinct architectures distributed throughout the SSN (Fig. S1B). Associated domains included those targeting sugars seen on human glycans, including potential fucosidases (GH141), β-galactosidases (GH16 (52) and GH165), mannosidases (GH92), sialidases (GH33) and O-acetyl esterases (CE4 and CE15) (Table 1). CE4 family members are deacetylases with substrates including amino sugars, though none characterized to date target Neu5Ac. CE15s are sparsely studied but so far are uniformly glucuronyl O-methylesterases.

**Table 1.** Domain arrangement of multifunctional GH156s.

| IDs | Architecture |
| --- | --- |
| GUT_GENOME011991_00921, GEM_PF-490474 | GH156 - GH141 |
| A0A1G1A1F0 | GH156 - GH16 |
| A0A1G3AKI5, A0A4V0XR61, GEM_PF-332583 | GH141 - GH156 |
| A0A2D5S979, A0A2E1KRC2, A0A2E7LA74, A0A2E8CZL7, A0A3L7UH35, GEM_PF-207205, GEM_PF-499370 | GH156 - GH165 |
| A0A356EQQ9 | GH33 - GH156 |
| GEM_PF-1731955 | GH156 - CE4 |
| GEM_PF-26113 | GH92 - GH156 |
| GEM_PF-354959 | GH156 - GH33 |
| GEM_PF-390790 | GH165 - GH156 |
| GEM_PF-598500 | GH156 - GH33 |
| GEM_PF-668270 | CE15 - GH156 |

### Genomes encoding GH156s also contain genes associated with degradation and metabolism of human glycans

We reasoned that organisms that use GH156s to break down host glycans would be able to release and/or utilize other sugar units composing these glycans. We therefore searched representative GH156-encoding UHGG genomes, predominantly in the form of MAGs, for genes related to the hydrolysis and metabolism of sugars found in human N- and O-glycans (Table S1, Fig. S2). Several factors limit the interpretability of these analyses. MAG completeness ranged from 52.72% to over 99% with a median of 90.53% (Table S2). Even MAGs with over 95% completeness may lack a quarter of the organism’s conserved core genes, such as those involved in sugar metabolism, and half of its variable accessory genes, such as those encoding GHs (83). Furthermore, homology-based annotation of protein functions by standard classifiers and default parameters is less reliable for distant relatives of commonly studied model organisms (i.e. Bacteroidota, Verrucomicrobiota and Firmicutes_A lineages), as the sequences have had greater opportunity to diverge. This has resulted in the functionally annotated proportion of a given proteome being highly dependent on taxonomic classification, with some genomes having as low as ∼20% of their proteome assigned functions (84). Despite these limitations, complete metabolic pathways can often be identified in MAGs and partial pathways can offer some evidence of functional capacity. To account for the incomplete nature of MAGs we grouped GH156-containing genomes at the taxonomic family level and evaluated the proportion of genomes from the family to encode a given function (Fig. 2). After removing families with fewer than 3 genomes, 2 Bacteroidota, 5 Firmicutes_A, and 3 Verrucomicrobiota families were analyzed.

**Figure 2.**
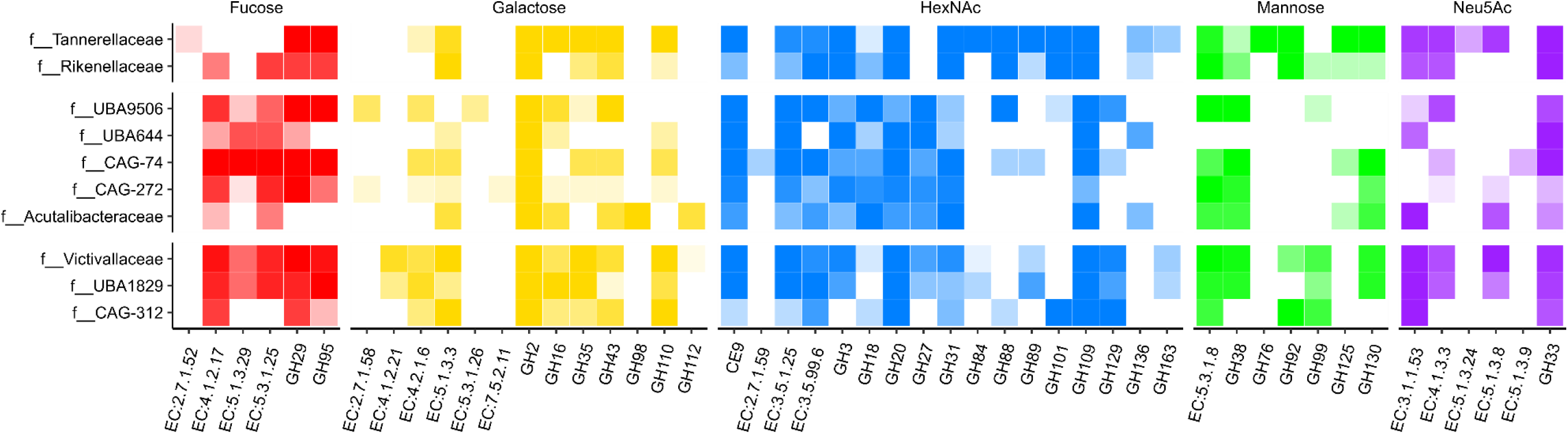
Genes involved in host-glycan monosaccharide hydrolysis and catabolism are frequently identified in GH156-encoding genomes. Genomes are grouped at the taxonomic family level and the proportion of genomes encoding a function is represented by opacity.

In general, GH156-encoding organisms appear to be capable of participating in N- and O-glycan catabolism by releasing and/or metabolizing the various monosaccharide constituents. Interestingly, all of these taxonomic families encode genes for GH33, which commonly display sialidase activity, potentially implying that these organisms exploit structurally distinct sialosides from diverse dietary and/or host sources. *N*-acetylneuraminate lyase (EC 4.1.3.3) catalyzes the first committed step in sialic acid catabolism. Its gene was identified in MAGs from 7 of the families across all three phyla and each of the families lacking a *N*-acetylneuraminate lyase gene encoded either sialate *O*-acetylesterase (EC 3.1.1.53) which liberates acetate from free or glycosidically-linked sialic acid, *N*-acylglucosamine 2-epimerase (EC 5.1.3.8) which participates in sialic acid and *N*-acetylmannosamine catabolism.

Genes encoding mannose-6-P isomerase (EC 5.1.3.8) were represented across all genomes, as well as at least one gene encoding a GH family member associated with mannose release from N-glycans (especially, GH92 and GH130), excepting f UBA644. Notably, none of the three f UBA644 genomes annotated exceeded 90% completeness (Table S2), providing a plausible explanation for the absence of genes linked to mannose metabolism.

*N*-acetyl-D-glucosamine/*N*-acetyl-D-galactosamine-6-phosphate deacetylase (3.5.1.25) and glucosamine/galactosamine-6-phosphate deaminase (3.5.99.6) play critical roles in GlcNAc and GalNAc degradation and genes encoding these functions were identified in all families, except for f UBA644. As well, several GH families were frequently observed across the genomes examined. Prominent examples include members of the exo-acting GH20 family, which are critical for HexNAc removal from host-glycans; the GH18 family, which includes endo-acting β-*N*-acetyl glucosaminidases active against the N-glycan chitobiose core, and; the α-GalNAc hydrolases GH109 and GH129, members of which have not been extensively studied, though representative substrates include the blood group A antigen and mucin GalNAc-α1-Ser/Thr (48).

The most common genes identified dedicated to galactose catabolism included those encoding aldose 1-epimerase (5.1.3.3) of the Leloir pathway and galactonate dehydratase (EC 4.2.1.6). Commonly observed galactosidase-containing GH families include GH110, members of which have been implicated in group B blood antigen hydrolysis. Both fucosidase GH families with members active on host glycans, GH29 and GH95, were frequently detected, as were L-fucose degradation I pathway constituents. The Tanerellaceae genomes were notably devoid of fucose-metabolizing genes, though they encoded GH29 and GH95, suggesting that these organisms might release fucose into the environment to the benefit of crossfeeders. For example, *B. thetaiotaomicron* cannot metabolize Neu5Ac, but uses sialidases to access the underlying glycan, the constituents of which it can utilize, and releasing Neu5Ac in the process (55).

### GH156-encoding genes are colocalized with genes encoding diverse GHs

In both Gram positive and Gram negative taxa, genes encoding GHs are often located on the chromosome alongside other enzymes, transporters and regulators involved in the saccharification of the same complex carbohydrate (27, 42). This polysaccharide utilization loci (PUL) architecture has been observed for the *B. thetaiotaomicron* breakdown of N- and O-glycans (85). We reasoned that colocalization of GH156 encoding genes with genes encoding GHs assigned to families known to contribute to host glycan degradation would provide support for the hypothesis that GH156s contribute to host sialoglycan degradation. To this end, we annotated the genes 20 kb up and downstream from the GH156-encoding genes using dbCAN2 (Fig. 3). While many GH156-encoding genes are colocalized with at least one other GH family gene potentially active against host glycans, none of the clusters examined display a complete complement of modules required for comprehensive degradation of a host glycan. Some Bacteroidota GH156s are encoded adjacent to and in the same orientation as GH92 genes. GH92s include α1-2, 3 and 6 mannosidases and *B. thetaiotaomicron* GH92s have been implicated in host N-glycan breakdown (55). Other Bacteroidota GH families include GH141 (L-fucosidases) and GH106 (L-rhamnosidases). Genes in proximity to Firmicutes_A GH156s encode putative galactosidases (GH2 and GH16), endo-β-*N*-acetylglucosaminidases (GH163), exo-α-*N*-acetylgalactosaminidases (GH109), sialidases (GH33), fucosidases (GH95), and a GH123, a family that includes a β-*N*-acetylgalactosaminidase active against host glycolipids. Notable GH families encoded alongside Verrucomicrobiota GH156 genes include exo-α-*N*-acetylgalactosaminidases (GH109), exo-α-galactosidases (GH110), and mannosidases (GH130).

**Figure 3.**
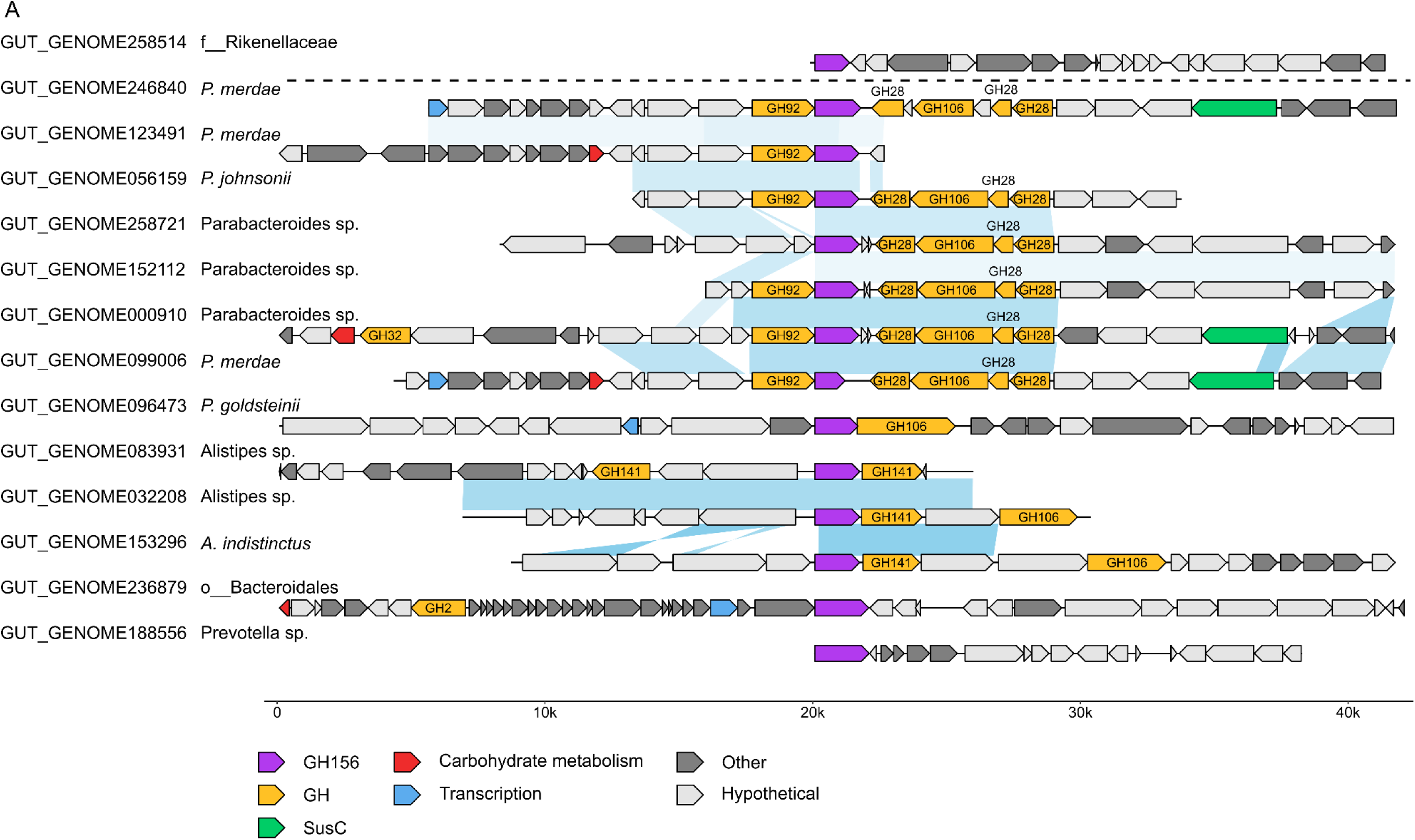
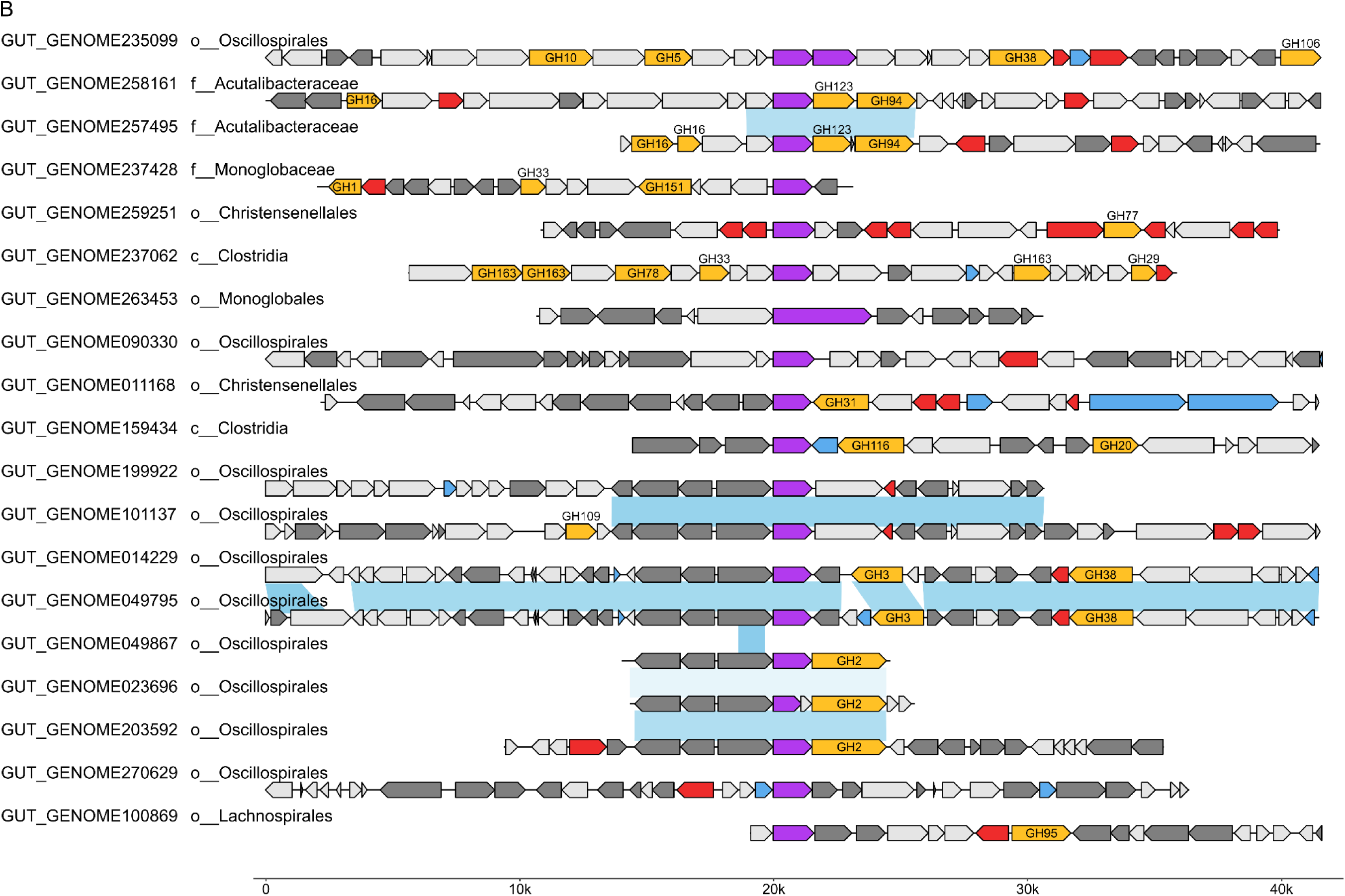
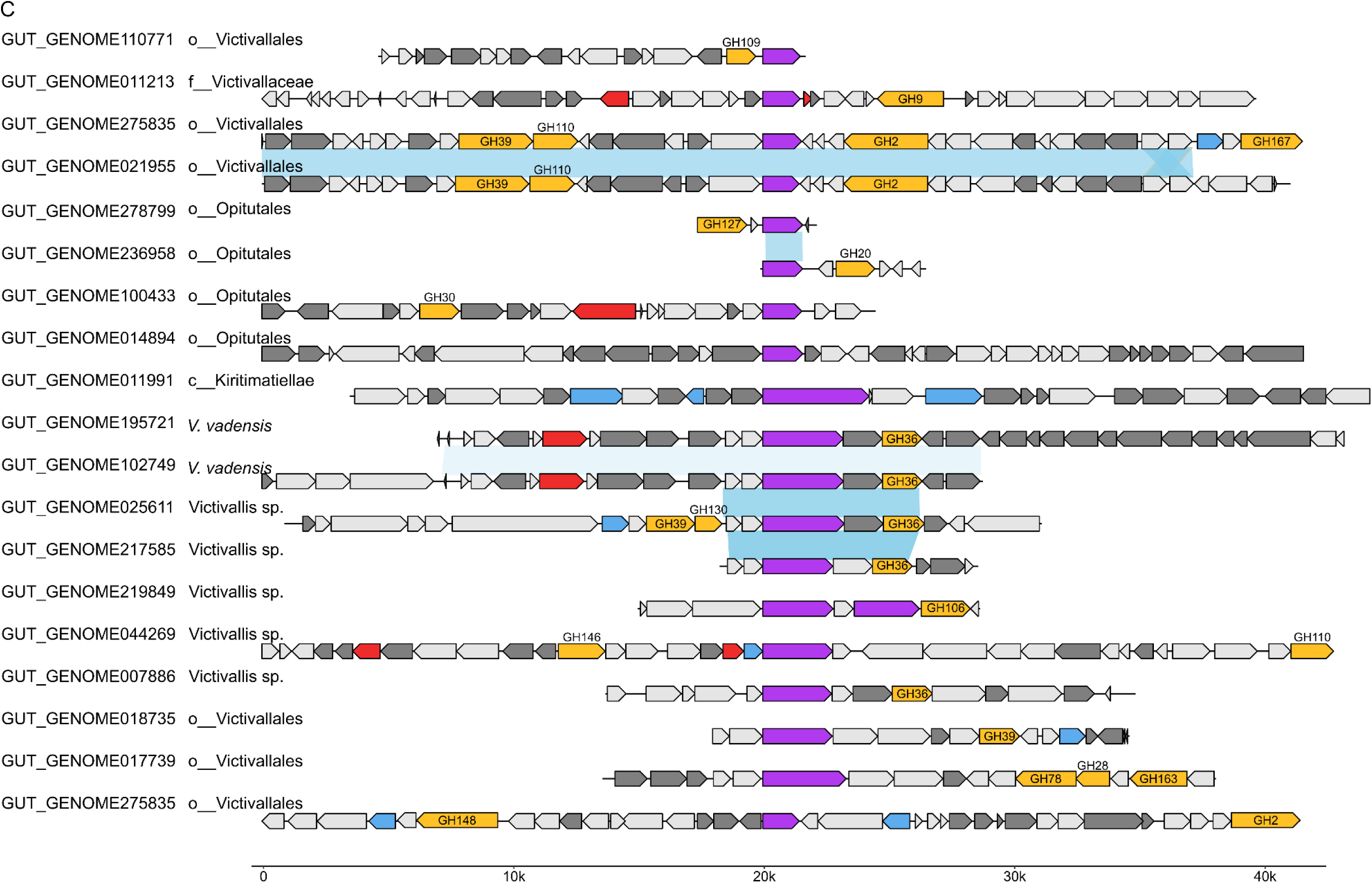
Genomic neighbourhood of GH156-encoding genes from the human gut microbiome. Organization of genes colocalized with UHGP-95-derived GH156-encoding genes from Bacteroidota (A), Firmicutes_A (B), and Verrucomicrobiota (C) genomes. Gene clusters below the dashed line in panel A contain genes encoding GH156s found in the same SSN cluster

### GH156 genes are more diverse and abundant in traditional populations than in industrialized Western populations

To investigate the distribution of the most common GH156 genes from the human microbiome, we used publicly available metagenomic data on distinct human populations. Reads from three studies (86–88) were mapped against GH156 domain nucleotide sequences. The proportion of bases covered for a given GH156 sequence with reads mapped from a single sample range from 1.5%-100% (Fig. 4A). A majority of sequences with very low coverage were derived from databases other than UHGP-95. Given that the reads were from gut metagenomes, we reasoned that most GH156s present in a sample would be from UHGP-95. Therefore, these low coverage genes were likely false positives, but some genes derived from non-gut databases with almost complete coverage were likely authentic and represent a gap in the UHGP-95 catalog. To help distinguish those genes likely present in a sample from those likely absent, at various coverage detection thresholds we calculated the proportion of detected sequences across all samples that were derived from UHGP-95 (Fig. 4B). We sought to maximize the proportion of UHGP-95-derived genes while minimizing the stringency of the threshold. At a coverage threshold of 275 bp (a local maximum), 96.4% of the GH156 sequences detected were from UHGP-95. We therefore filtered our data to remove hits with less than 275 bp of coverage in each sample. The two remaining non-UHGP-95 genes detected in at least one sample included PWL98936.1 from Uniprot and GEM-PF_16266 from GEM, which are both derived from human fecal metagenomes, providing strong support that all hits included in the analyses are truly present in the sample.

**Figure 4.**
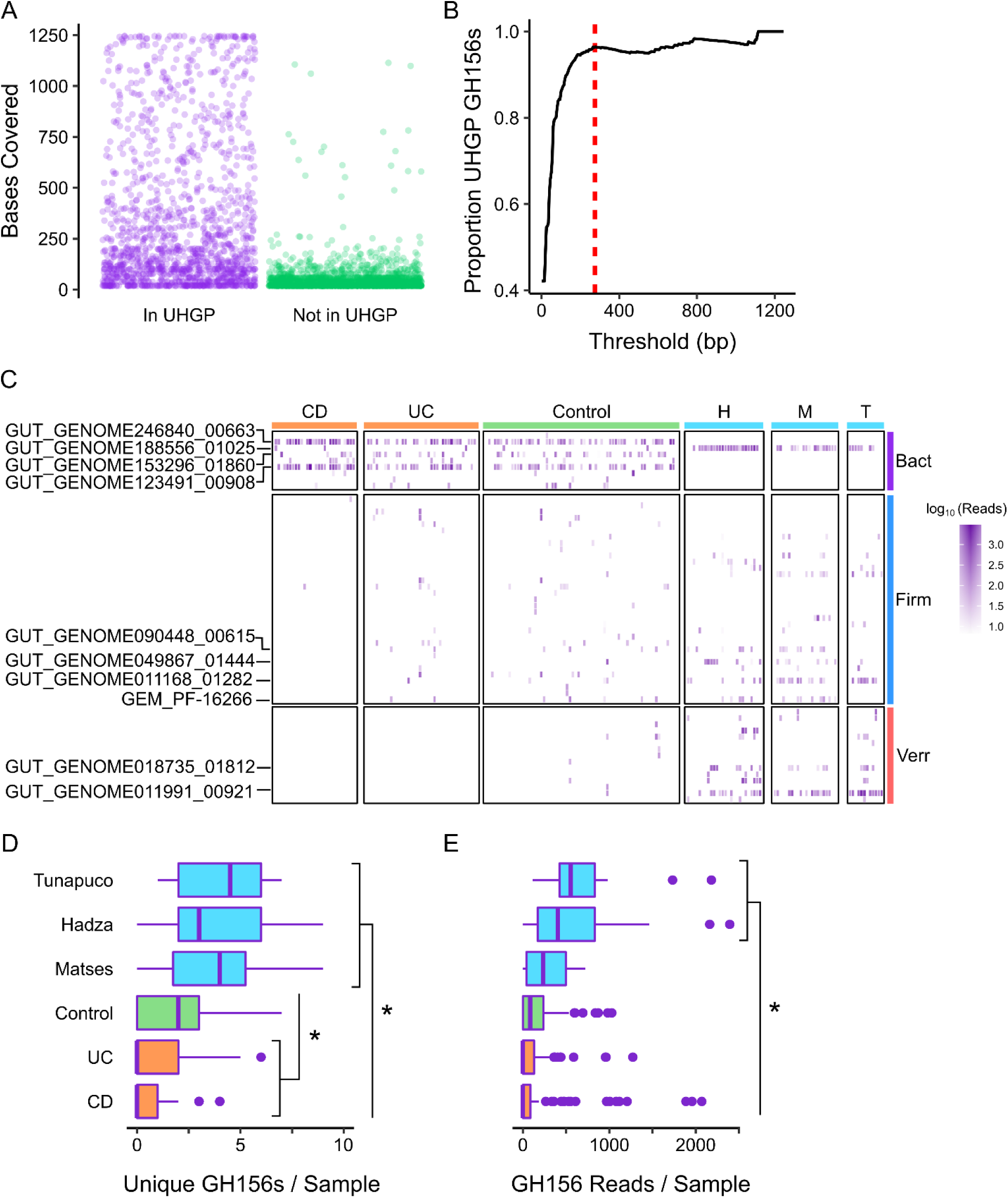
A greater variety and abundance of GH156s are associated with the metagenomes of individuals practicing traditional lifestyles relative to individuals from industrialized societies or with IBD. Raw reads from metagenomic datasets were mapped to GH156-encoding genes (A). Proportion of genes detected across all samples derived from UHGP-95, depending on the coverage threshold used for detection (B). GH156s detected in each sample and their relative abundance (C): H, Hadza; M, Matses; T, Tuanpuco; Bact, Bacteroidota; Firm, Firmicutes_A; Verr, Verrucombicrobiota. The ten most prevalent GH156s are labelled. Number of distinct GH156s detected in each sample (D) and number of reads mapped to a GH156 gene in each sample (E). The * denotes p value less than 0.05.

Metagenomic reads were obtained from IBD (CD or UC) patients or non-IBD controls from industrialized societies (Italy, Oklahoma, Massachusetts) as well as from traditional hunter-gatherer (Hadza, Matses) or agriculturalist communities (Tunapuco) (Fig. 4C). Relative to Western industrialized societies, traditional lifestyles are typically correlated with a wider variety and greater abundance of dietary complex carbohydrates (from wild plants) (87–89) and are believed to be associated with lower incidences of IBD (90). The Hadza metagenome encodes a correspondingly greater variety and abundance of CAZymes relative to industrialized populations. Consistent with this, traditional foragers and agriculturalists metagenomes contained a greater number of unique GH156 encoding genes (Fig. 4D). In turn, individuals with IBD had fewer than healthy western controls. The assortment of GH156s found in traditionalist metagenomes was distinct from those of western individuals (Fig. 4C). While GH156 genes from Verrucomicrobiota are absent from CD and UC patients and most healthy controls, they are prevalent in the hunter-gatherer/farmer groups. The Firmicutes_A GH156 genes are also seen more frequently in the traditional lifestyle groups than in the IBD groups. Consistent with typical IBD dysbiosis of reduction in Firmicutes and Verrucomicrobiota and expansion of facultative anaerobes and Actinobacteria (91), which do not encode GH156 genes. Conversely, a single Bacteroidota GH156 from a *Prevotella* species is detected in traditional forager/agriculturalist populations, while the post-industrialized groups largely contain GH156s from a mix of *Parabacteroides* species. The ‘Bacteroidota’ cluster (Fig. 1D, red circle) contains the four most frequently detected GH156s (Table S3). This includes the aforementioned *Prevotella* GH156 as well as the GH156 from *P. merdae,* which is one of 57 species in >90% human gut microbiomes sampled globally (92). The abundance of a given GH156 gene in a sample is estimated by the number of reads mapped to that gene. Hadza and Tunapaco metagenomes contained a greater abundance of GH156-encoding genes than the other groups examined (Fig. 4E), consistent with the prevalence observed for other GH families (87–89).

### The GUT_GENOME011168_01282 protein exhibits sialidase activity

To facilitate molecular cloning of genes encoding GH156s from the human intestinal tract we mapped metagenomic raw reads from healthy volunteer fecal samples to the GH156 gene sequences as previously described. Two genes encoding GH156s from the same SSN cluster as EnvSia156 (Fig. 1D) could be detected (data not shown): GUT_GENOME011168_01282 (d Bacteria;p Firmicutes_A;c Clostridia;o Christensenellales;f CAG-74;g ;s) and GUT_GENOME014894_00745

(d Bacteria;p Verrucomicrobiota;c Verrucomicrobiae;o Opitutales;f CAG-312;g CAG -312;s CAG-312 sp000438015). Notably, GUT_GENOME011168_01282 was the sixth most frequently observed GH156 across all metagenomes examined, predominantly in subjects practicing traditional lifestyles but also in three healthy controls and one UC patient (Table S3). Global alignment of EnvSia156, and a second GH156 with confirmed sialidase activity (OIO94155.1) (69) with the gut derived GH156 sequences detected showed relatively low sequence identity, in the range of 27-35% (Fig. 5A). The two gut-derived GH156 genes were cloned from DNA obtained from feces into IPTG-inducible expression vectors to facilitate recombinant polyhistidine-tagged protein production. Batch purification of GUT_GENOME014894_00745 and GUT_GENOME011168_01282 using Ni^2+^-NTA resin resulted in similar yield and purity, as judged by SDS-PAGE (Fig. 5B). To evaluate sialidase activity, the recombinant GH156s were incubated with 100 μM of the fluorogenic substrate 2’-(4-methylumbelliferyl)-α-D-*N*-acetylneuraminic acid (4MU-Neu5Ac) for 30 minutes at 37 °C. Hydrolysis of 4MU-Neu5Ac results in an increase in fluorescent signal which can be detected spectrofluorometrically. A significant increase in fluorescence relative to the control suggests that GUT_GENOME011168_01282 is a sialidase (Fig. 5C). Unfortunately, GUT_GENOME014894_00745 did not liberate Neu5Ac from 4MU under any of the conditions tested.

**Figure 5.**
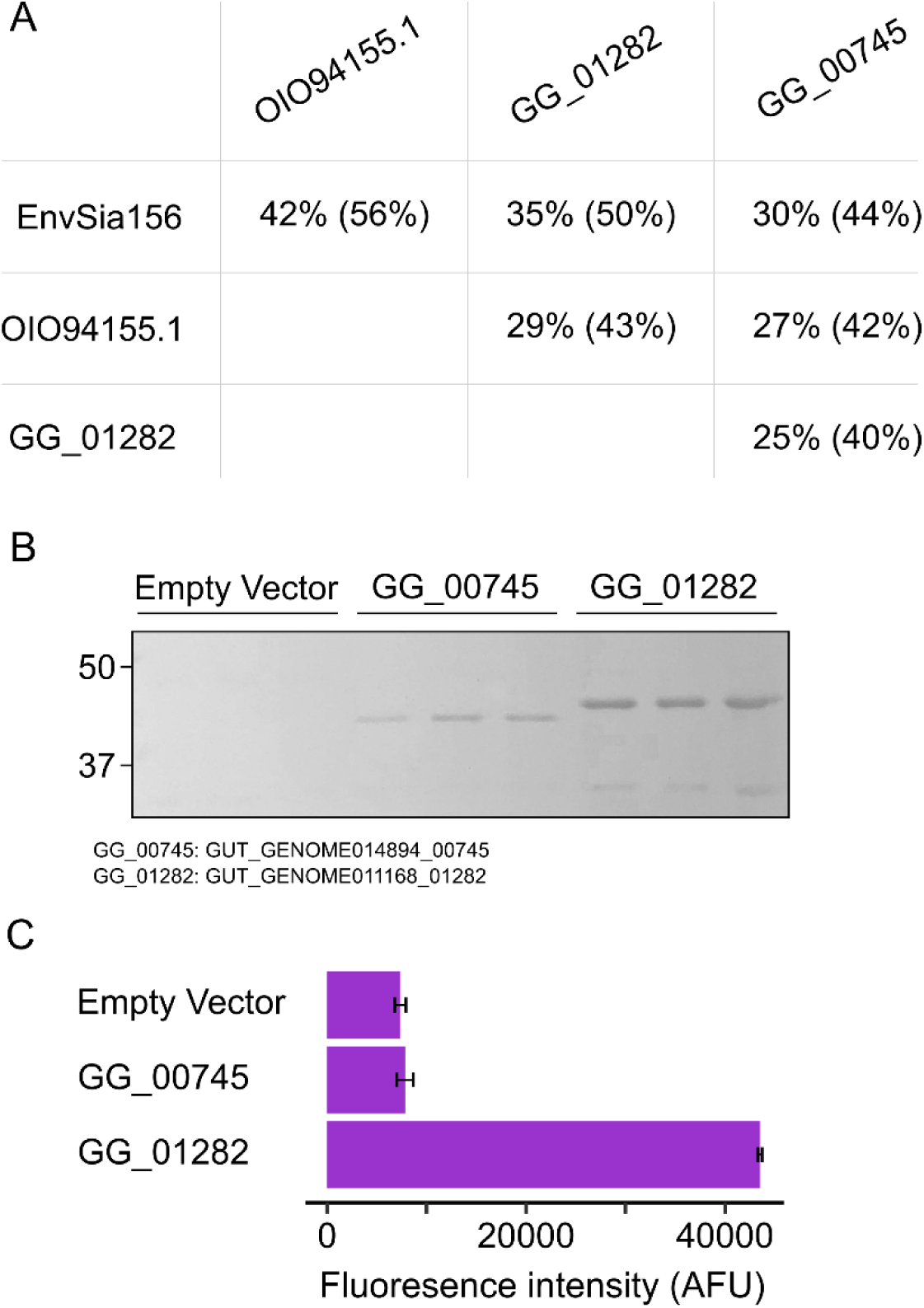
A GH156 prevalent in the gut metagenomes of individuals from traditional societies displays sialidase activity. Pairwise percent identity (percent similarity) of GH156s with confirmed sialidase activity and candidate sialidases (A). Poly-histidine-tagged recombinant GH156s from gut metagenomes were batch purified (B). A fluorescence-based end-point assay using recombinant enzymes and 4MU-Neu5Ac as a substrate revealed GUT_GENOME011168_01282 is a sialidase (C). Error bars indicate standard error across biological triplicates.

## Discussion

GHs active on host-produced glycans play important roles in the gut environment, and impact microbiota community structure and virulence. This is particularly true for sialidases, since sialic acids occupy terminal positions and can act as receptors for microorganisms and immune cells, are an abundant and accessible nutrient source, and contribute to the physiochemical properties of mucous. To this end, we are interested in characterizing novel sialidases. We used a structure-informed homology search of large protein databases to search for members of the newly described GH156 family. At the time of writing, the CAZy database catalogues 51 members of this family, only two of which has demonstrated sialidase activity. By including protein sequences from metagenome datasets in our search, we were able to identify ∼10-fold more putative family members that can plausibly act on sialic acid, given homology to EnvSia156 and the presence of critical Neu5Ac-binding and catalytic residues. This includes 90 unique protein sequence from human fecal samples.

The proposed GH156s are largely derived from poorly studied phyla, including Planctomycetes, Verrucomicrobiota, Firmicutes_A and Bacteroidota (Fig. 1B), a potential indication as to why it was the ∼160^th^ GH family described. These taxa remain difficult to isolate, indeed *P. merdae, P. goldsteinii*, *P. johnsonii*, *Alistipes_A indistinctus*, and *V. vadensis* were the only isolate whole genome sequences to contain a GH156, and the identification of so many GH156s was only possible due to deep sequencing and MAG assembly. The tremendous abundance of GH genes in Bacteroidota genomes is well established, and has been noted for the Verrucomicrobiota gut species *Akkermansia muciniphila* and *V. vadensis* (27, 43). Planctomycetes are important degraders of complex carbohydrates in a great variety of soil and aquatic environments, both extreme and otherwise, suggesting many of the identified GH156s likely target plant glycans (93). Comparative genomics studies of Planctomycetes have revealed these genomes often contain dozens of genes encoding close and distant homologs of CAZymes, suggesting this phylum may be an abundant repository of novel GH families with activities that evolved for their varied niches (94). The GH156 sequences we identified were quite heterogeneous (Fig. 1DE), likely because only a small fraction of extant sequences have been identified and deposited in public databases. Assembly of genomes of gut microbes from undersampled human populations (Fig. 1C) as well as samples of other animals and environments, will go a long way in filling in the gaps in order to build a more informative SSN with monofunctional clusters (56, 57).

A possible implication of the sequence diversity of GH156s identified is a commensurate intra-family substrate diversity. This is consistent with the taxonomic profile of this family, suggesting that GH156s are active against complex carbohydrates present in various environments. Though it is generally used to refer to Neu5Ac, ‘Sialic acid’ is a generic name given to a large group of nine-carbon acidic sugars (nonulosonic acids), any of which could potentially be targeted by a given GH156 (95, 96). Common examples of sialic acids include N-glycolylneuraminic acid (Neu5Gc), 2-keto-3-deoxy-D-glycero-D-galacto-nononic acid (Kdn), Pseudominic acid and Legionminic acid. Smaller ulosonic acids, including 3-deoxy-D-manno-oct-2-ulosonic acid (Kdo) and 2-keto-3-deoxy-D-lyxo-heptulosaric acid (Dha), are additional candidate substrates and GH33 family members target these sugars as well (97). Indeed, the EnvSia156 residues in proximity to the Neu5Ac acyclic functional groups are variable amongst family members (Fig. 1G). Though the structure of the founding member shows the C5 position extends away from the enzyme, rather than in a hydrophobic pocket. The sialic acid acyclic glycerol group (C7-C9) makes extensive contact with the binding site. Together, its likely that EnvSia156 homologs have binding-site architectures that accommodate different ulosonic acid substrates. Unfortunately, fluorogenic substrates are not readily available to test these various activities, complicating biochemical assays.

Generally, organisms that encode GH156 genes also contain genes that may be involved in the liberation and catabolism of a variety of sugars that compose human N- and O-glycans (Fig. 2). This offers some indirect evidence that the organisms could be mucous-degraders, though direct experimental evidence is required to resolve this. Assigning glycan substrates is especially fraught given the polyfunctionality of well described CAZy families (e.g. GH16), and the poor characterization of other families with critical mucolytic activities (e.g. GH101 and GH129). Unfortunately, genomic investigation at the level of the GH156 gene cluster was not able to provide more definitive insights into the matter, by ‘guilt-by-association’ with a set of families known to act on host-glycans (Fig. 3). Interestingly, GH156s were often associated with GH141s; either as separate proteins encoded in proximity to GH156 genetic loci or as separate domains in the same GH156 protein. One of the two GH141s that has been studied displayed fucosidase activity on a plant cell wall substrate, but its possible that GH141s are also active against host fucoglycans. Furthermore, gaps in sugar catabolic pathways could be due in part to the phyla under investigation being poorly described and only distantly related to model organisms in which the pathway functions were originally discerned. Interestingly, many organisms that carry a GH156 gene also carry a GH33 gene. These genes may be redundant, recognize different ulosonic acid substrates, or recognize the same sugar in different macromolecular contexts (e.g. linkage type or specific aglycone). A reverse genetics approach monitoring upregulation of GH156s in isolates growing on host-glycans can help resolve glycan substrates (55, 97).

Although relative to Firmicutes_A and Verrucomicrobiota, Bacteroidota encode few discrete GH156s (Fig. 1B), these enzymes are the most prevalent in the gut metagenomes examined (Fig. 4C). Notably, only Bacteroidota GH156s could be detected in multiple IBD samples, and Verrucomicrobiota GH156s were rarely detected. Relative to samples from industrialized societies, the Hadza, Metses and Tunapuco gut metagenomes encoded a larger variety of GH156s from Firmicutes_A and Verrucomicrobiota taxa and a single Bacteroidota GH156 from an unclassified *Prevotella* species. Traditional lifestyles are often associated with greater gut microbiota diversity and this has been credited to the consumption of a greater variety and abundance of plant material. In this light, it seems likely that the GH156s harboured in their microbiomes would contribute to plant fibre degradation or the organisms are generalists that can switch between host and dietary glycans. Here we were able to demonstrate the Neu5Ac sialidase activity of a GH156 commonly encoded in the gut metagenomes of individuals living traditional lifestyles.

*In vivo* targeting of Neu5Ac is a promising avenue for cancer therapies (98, 99). Cancer cells often hypersialate the cell surface to evade the immune system via Siglec-binding. Bacterial sialidase-antibody conjugates have shown efficacy in proof-of-concept experiments making them promising therapeutics (100). Development of this technology involved screening a number of different GH33 sialidases to choose one with optimal activity against host glycans, demonstrating a need to catalogue the activities of representative sequences across the sialidase sequence space is critical for identifying enzymes of biotechnological importance. The GH156s identified here offer an additional pool of candidates with potential host sialoglycan activity. Of particular interest are the GH156s from the from the human gut, since there is an increased probability they would be active against host sialic acids relative to candidates from other environments. This rationale lead to the discovery of blood group specific GHs from the human gut using a functional metagenomics screen (101). Functional screens using random fragment libraries and rational selection of putative GH sequences from genomes and metagenomes are both important strategies for uncovering functional novelty (102). The GH156 sequence space remains almost entirely unexplored and promises to offer new biochemical activities.

## Materials and Methods

### Bioinformatics methods

HmmerBuild (v3.1b2) (71) was used to construct a profile Hidden Markov Model using default settings and a multiple sequence alignment made using MAFFT (v7.471) G-INS-I (103). Skylign webserver was used to construct the sequence logo (104). HmmerSearch was used for database searching. CD-HIT (v4.8.1) was used with default settings and a sequence identity cut-off threshold of 0.95 to remove redundant sequences (75, 76). Sequence Similarity Networks were calculated using the SSN tool developed by the Enzyme Function Initiative (77–80), with a supplied multi-fasta file and an initial alignment score of 10. Edges were filtered and the SSN visualized using Cytoscape (v3.8.2) (105). To depict sequence conservation, the EnvSia156 6S00 structure was coloured by sequence conservation using Chimera v1.14 (106). The sequence alignment to facilitate this was generated by predicting the 3-dimensional structure of 79 UHGP-95 GH156s using Alphafold2 with ColabFold MMseqs2 v1.3 (107, 108) and generating a multiple structural alignment along with pdb 6S00 using mTM-align (109).

Publicly available metagenome datasets PRJNA400072 (88 CD, 76 UC, 56 Control) (86), PRJNA268964 (24 Matses, 12 Tunapuco, 22 control) (88) and PRJNA278393 (27 Hazda, 11 control) (87) were downloaded from the Sequence Read Archive (SRA) using entrez direct. We then mapped raw reads to GH156 genes truncated to the GH156 domain boundaries using bwa mem (110). Mapped reads were sorted using Samtools view (111), and the number of mapped reads were identified via Samtools coverage. Read counts were adjusted to account for variability in gene length and per sample read depth using the formula: number of reads × (maximum GH156 gene length / mapped GH156 gene length) × (maximum number of reads in all samples / number of reads in sample). Samples with adjusted read counts more than 3 standard deviations above the mean were removed from the analysis.

GH156 accessory domains were predicted using the dbCAN2 metaserver (82). UHGG genomes (Table S2) were annotated with DRAM (112) to retrieve EC numbers (Table S1), and with the dbCAN2 metaserver to assign CAZyme families.

Genes on the same contig 20 kb up or downstream of GH156 were searched for CAZymes using the dbCAN2 meta server. Clusters were visualized with the R package gggenomes v.0.9.5.9 (https://github.com/thackl/gggenomes). The minimap2 software was used to detect synteny (113).

### Statistical methods

Statistical analysis and visualization was performed with R v.4.0.3 (114) with the tidyverse package (115). Significant differences were determined using the rstatix package v.0.5.0 Tukey Honest Significant Differences (TukeyHSD) function.

### Bacterial strains and growth conditions

Bacterial cultures were grown at 37 °C with aeration in lysogeny broth (LB) supplemented with 50 μg ml^-1^ kanamycin. *E. coli* XL1-Blue (*recA1 endA1 gyrA96 thi-1 hsdR17 supE44 relA1 lac* [F′ *proAB lacI*^q^*ZΔM15* Tn*10* (*Tet*^r^)]) (Stratagene) was used for general cloning and *E. coli* BL21-CodonPlus(DE3) (F^-^ *ompT hsdS*(rB^-^ mB^-^) *dcm*^+^ *Tet*^r^ *gal* λ(DE3) *endA* Hte [*argU proL Cam*^r^] [*argU ileY leuW Strep/Spec*^r^]) (Stratagene) was used for recombinant protein expression.

### DNA methods

Custom oligonucleotide primers were obtained from Integrated DNA Technologies. PCR amplification was performed using Phusion Polymerase (New England Biolabs). Template material was metagenomic DNA from feces. The gene for GUT_GENOME011168_01282 was amplified using primers with the sequences 5′- gatc**gctagc**ATGCGACAGGGCATCATC-3′ and 5′-gatc**aagctt**TTACTCGAATACGCGCGCG-3′ and the gene for GUT_GENOME014894_00745 was amplified using primers with the sequences 5′-gatc**gctagc**TTTTTTGAAAAAATTTTTTTCTATTG-3′ and 5′- gatc**ctcgag**TCAGCCGGCATCGACAAAAATC-3′ (uppercase bases are homologous to the region being amplified and bolded bases are restriction enzyme recognition sites). The Monarch DNA Gel Extraction and Monarch PCR and DNA Cleanup kits (New England Biolabs) were used to clean PCR products according to manufacturers instructions. PCR amplicons were ligated into the kanamycin resistant vector pET28b(+) (Novagen) using T4 ligase (New England Biolabs) following restriction endonuclease digestions (NheI, XhoI, and HindIII; New England Biolabs) and PCR cleanup.

### Protein expression and purification

Protein was expressed and purified in three parallel batches from separate cultures. In 5 mL tubes, BL21-CodonPlus(DE3) harbouring expression plasmids were grown in LB to an A_600 nm_ of 0.6. Cultures were then incubated with aeration at 15 °C for 30 minutes prior to induction with 0.1 mM IPTG. Cultures were incubated for 20 hours at 15 °C with aeration, and a culture volume of 3 mL A_600 nm_ ^-1^ was harvested by centrifugation at 4,000 x g. Cells were lysed with BugBuster (Millipore) containing rLysozyme (Millipore) and Benzonase nuclease (Millipore) for 1 hour at room temperature on a rocking platform. Lysates were subject to centrifugation for 20 minutes at 20,000 x g to pellet intact cells and insoluble material and the supernatant was incubated with HisPur Ni-NTA resin (Thermo Fisher Scientific) equilibrated with buffer A [25 mM Tris, pH 7.4, 150 mM NaCl] + 25 mM imidazole at room temperature for 1 hour on a rocking platform. The resin was collected by centrifugation, washed 3 times in buffer A + 50 mM imidazole, and eluted with buffer A + 250 mM imidazole in a final volume of 150 μL. Eluted protein was exchanged into buffer B [50 mM NaOAc, pH 5.2, 100 mM NaCl] using drop dialysis with 0.025 μm cellulose filters (Millipore). Protein expression and purity were monitored using SDS-PAGE with 12% gels and Tris-glycine buffer (116), stained with SimplyBlue SafeStain (Invitrogen).

### Sialidase assay

Sialidase activity was fluorometrically monitored using the substrate 2’-(4-methylumbelliferyl)-α-D-*N*-acetylneuraminic acid (4MU-Neu5Ac; Sigma-Aldrich). Reactions were performed in a final volume of 50 μL. To 45 μL batch purified protein, 4MU-Neu5Ac was added at a final concentration of 100 μM in buffer B. The reaction was incubated at 37 °C for 30 minutes, and fluorescence intensity was measured with a BioTek Synergy H1 microplate reader (excitation: 365 nm, emission: 445 nm). Assays were performed in black-walled 96 well plates. and technical duplicates.

## Acknowledgements

EM is supported by a Postdoctoral Fellowship from the Canadian Institutes of Health Research. MGS is supported by a Canada Research Chair. This research was funded by a Proof of Principle grant from the W. Garfield Weston Foundation to EM and MGS, and a Canadian Institutes of Health Research grant to MGS. The authors declare no conflicts of interest.

---

**Figure S1.**
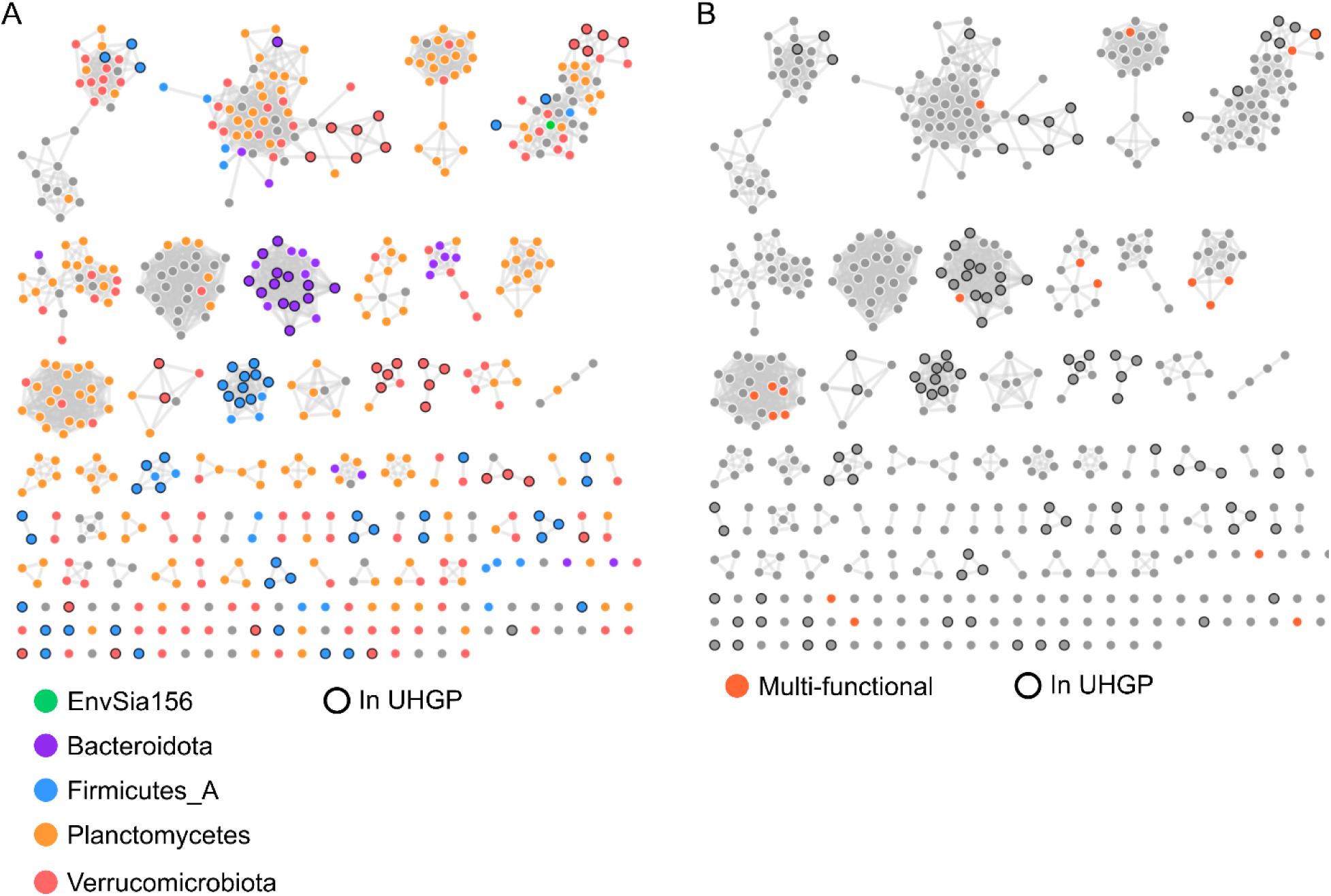
Taxonomic and modular distribution of GH156s in the family sequence space. SSN of GH156 domains with edges indicating >45% identity colored by phylum (A) or by presence of a second catalytic domain (‘Multi-functional’) in the full-length sequence (B).

**Figure S2.**
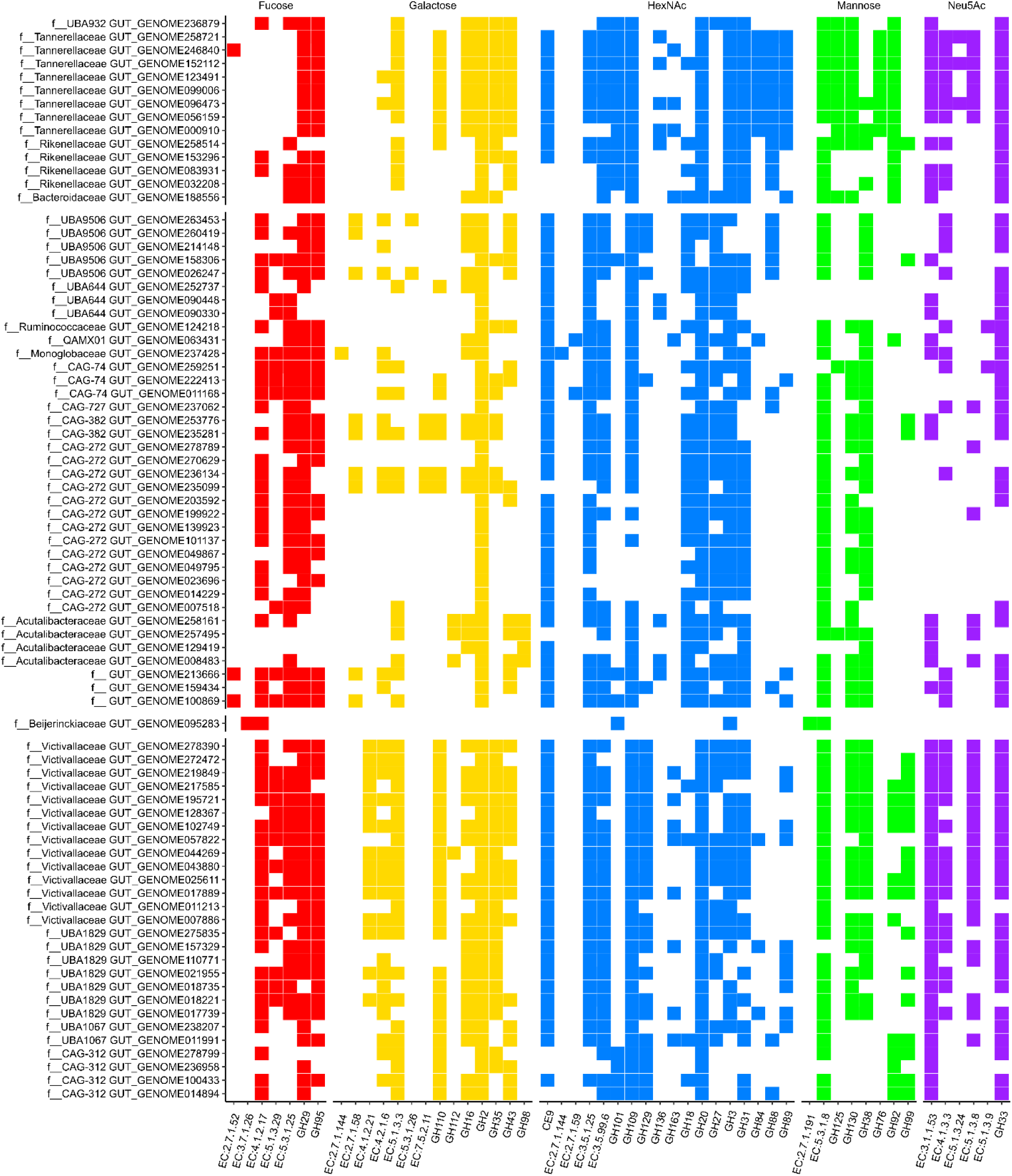
Presence of host-glycan sugar liberating or metabolizing genes in representative UHGG MAGs containing a GH156-encoding gene.

**Table S1.**
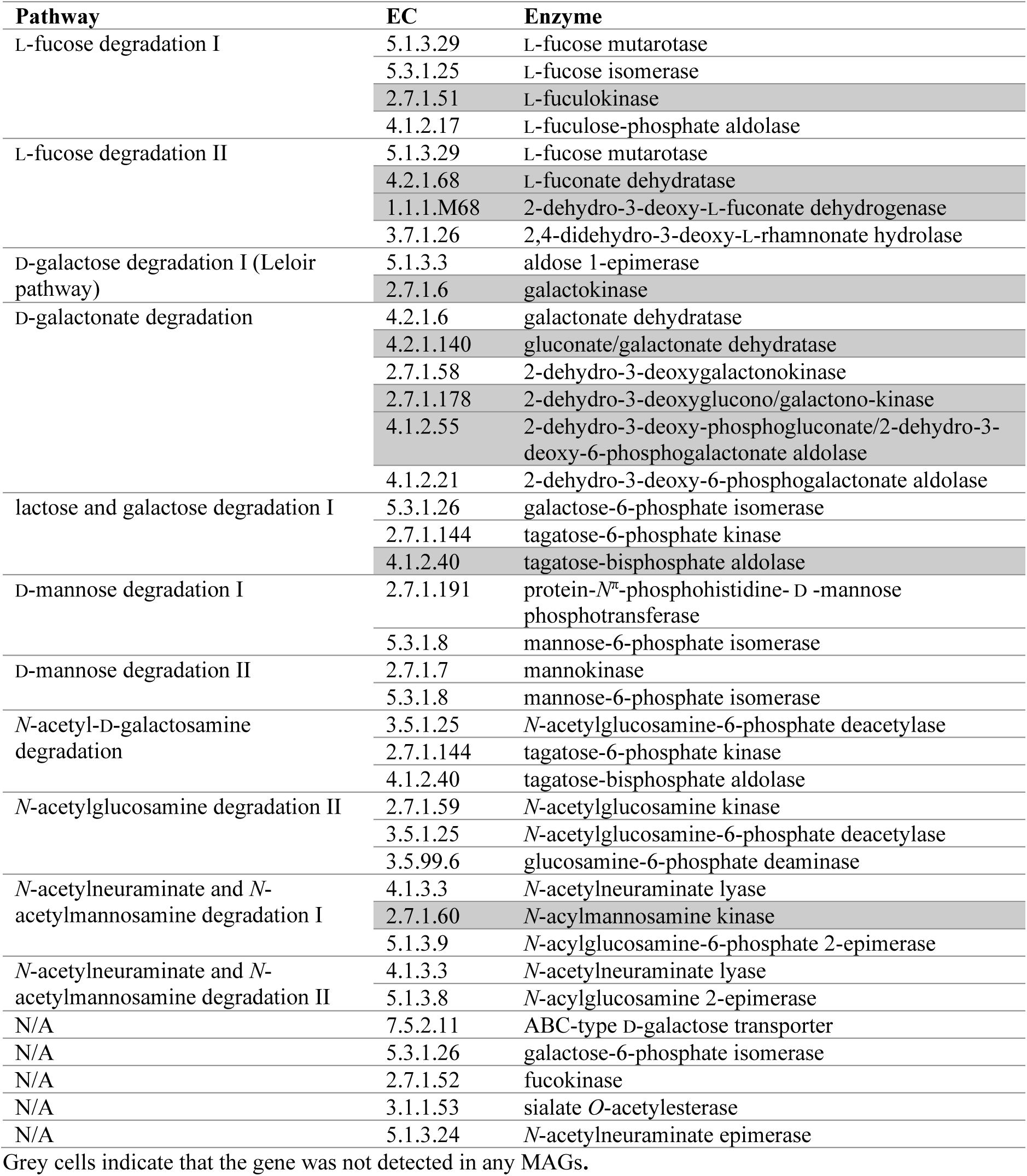
Simple sugar metabolic pathways and EC numbers searched for in MAGs.

**Table S2.**
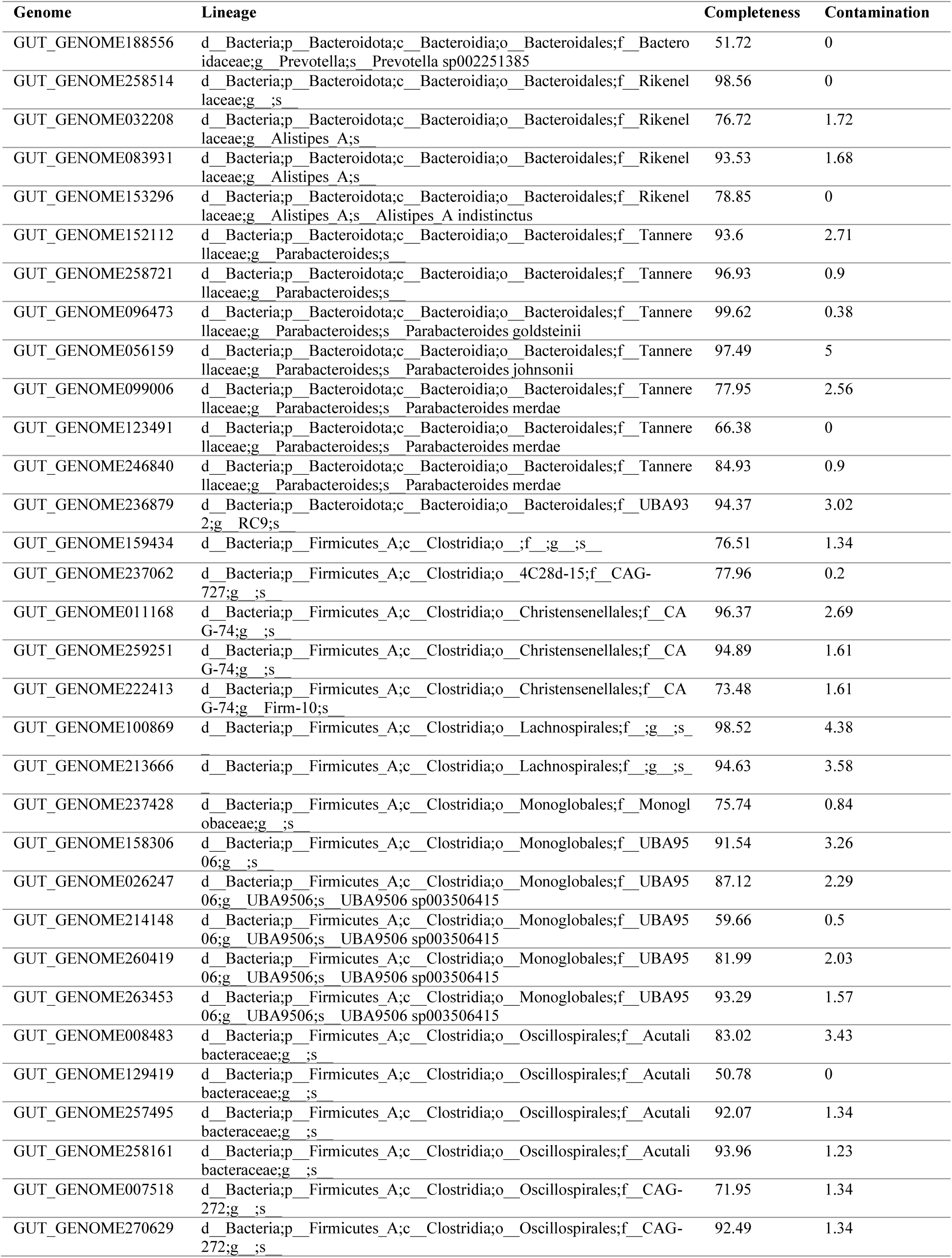
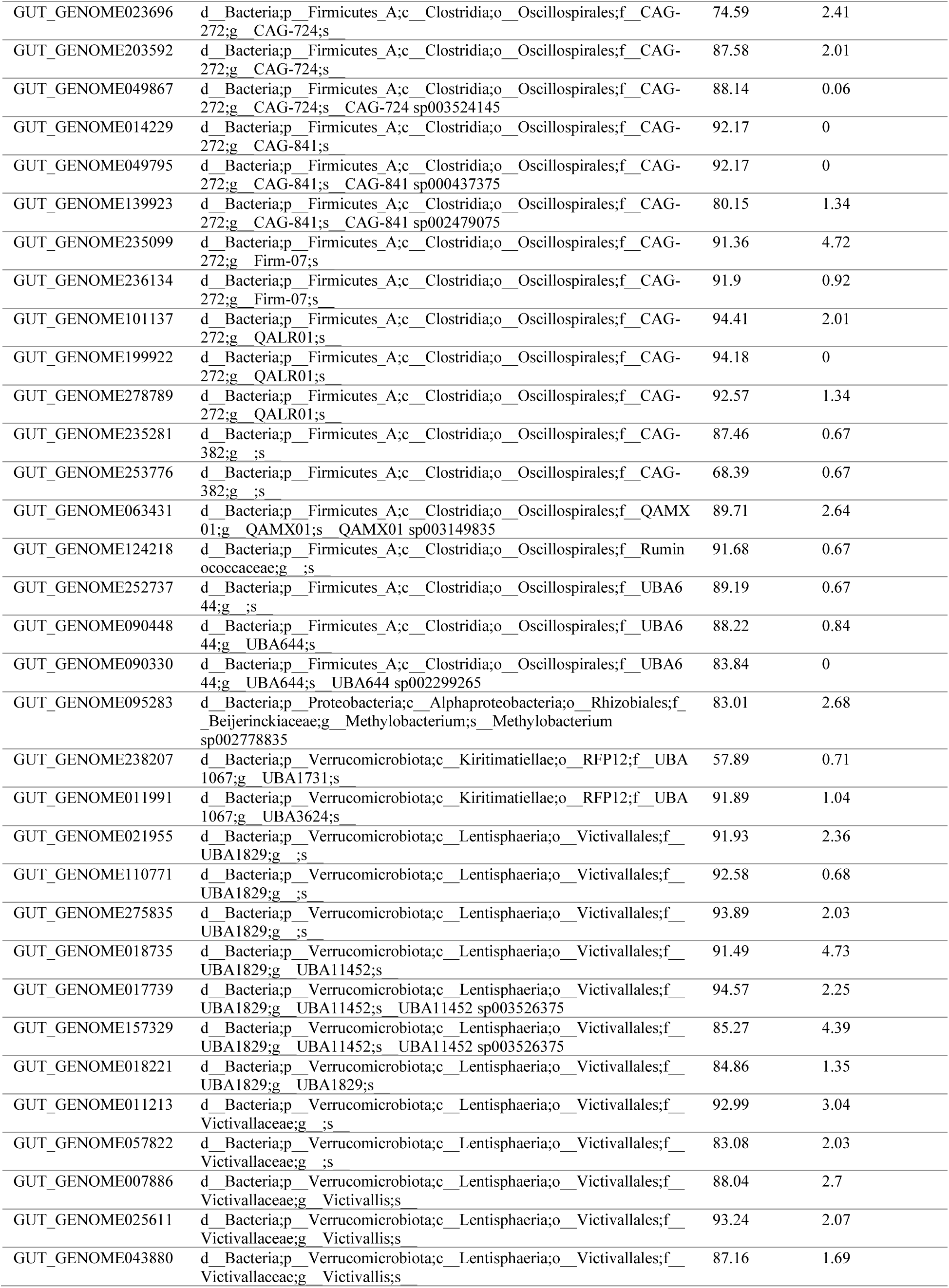
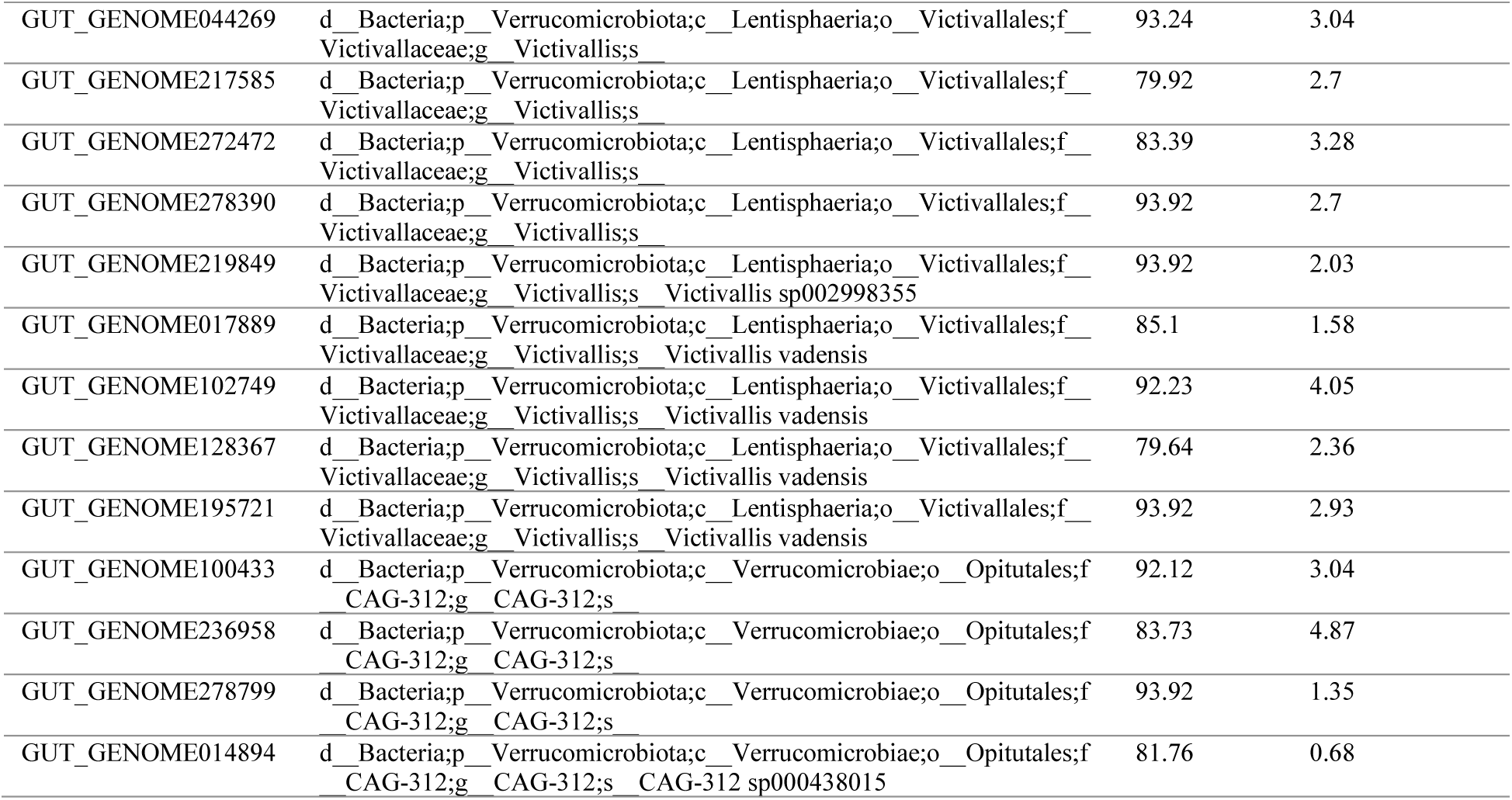
Completeness and contamination of UHGG MAGs containing genes encoding GH156s.

**Table S3.**
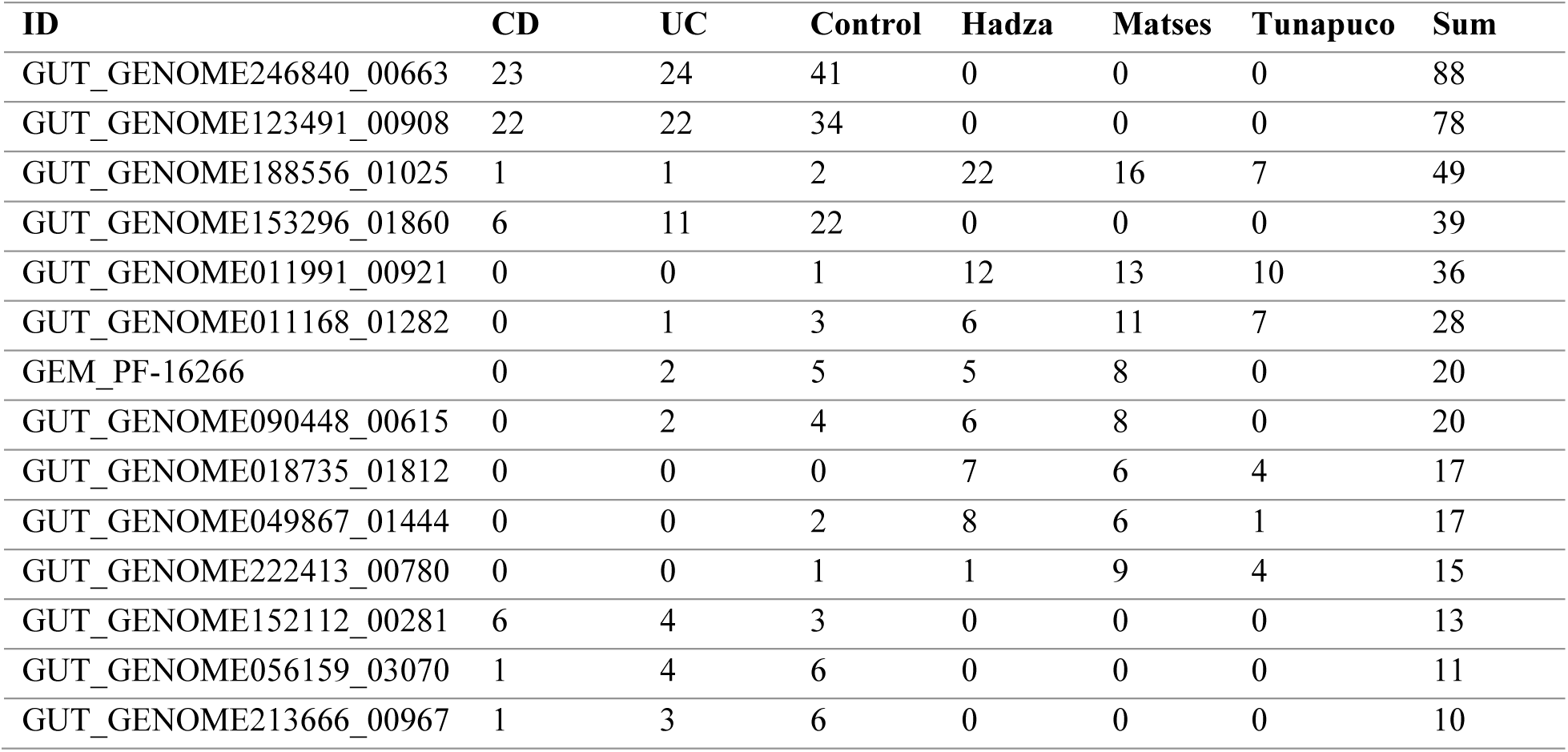
Most commonly detected GH156-encoding genes.

| ID | CD | UC | Control | Hadza | Matses | Tunapuco | Sum |
| --- | --- | --- | --- | --- | --- | --- | --- |
| GUT_GENOME246840_00663 | 23 | 24 | 41 | 0 | 0 | 0 | 88 |
| GUT_GENOME123491_00908 | 22 | 22 | 34 | 0 | 0 | 0 | 78 |
| GUT_GENOME188556_01025 | 1 | 1 | 2 | 22 | 16 | 7 | 49 |
| GUT_GENOME153296_01860 | 6 | 11 | 22 | 0 | 0 | 0 | 39 |
| GUT_GENOME011991_00921 | 0 | 0 | 1 | 12 | 13 | 10 | 36 |
| GUT_GENOME011168_01282 | 0 | 1 | 3 | 6 | 11 | 7 | 28 |
| GEM_PF-16266 | 0 | 2 | 5 | 5 | 8 | 0 | 20 |
| GUT_GENOME090448_00615 | 0 | 2 | 4 | 6 | 8 | 0 | 20 |
| GUT_GENOME018735_01812 | 0 | 0 | 0 | 7 | 6 | 4 | 17 |
| GUT_GENOME049867_01444 | 0 | 0 | 2 | 8 | 6 | 1 | 17 |
| GUT_GENOME222413_00780 | 0 | 0 | 1 | 1 | 9 | 4 | 15 |
| GUT_GENOME152112_00281 | 6 | 4 | 3 | 0 | 0 | 0 | 13 |
| GUT_GENOME056159_03070 | 1 | 4 | 6 | 0 | 0 | 0 | 11 |
| GUT_GENOME213666_00967 | 1 | 3 | 6 | 0 | 0 | 0 | 10 |

